# Physicochemical and functional comparison of food-grade and precision-fermented bovine lactoferrin

**DOI:** 10.1101/2025.03.03.641326

**Authors:** Jonathan Cavallo, Jared Raynes, Samuel Mandacaru, Deepa Agarwal, Lloyd Condict, Stefan Kasapis

## Abstract

This study provided a comprehensive physiochemical and functional comparison between food-grade, commercially available bovine lactoferrin (cLF) and two precision-fermented variants (pfLF_highFe and pfLF_lowFe) at varying iron saturation levels. LC-MS and FTIR/CD spectroscopy confirmed high purity and structural fidelity at primary and secondary levels, while UV-vis analysis revealed comparable molar absorptivity across all preparations. Fluorescence spectroscopy demonstrated tertiary structural variability with redshifts observed as a result of iron binding. Molecular weight (M_w_) comparisons made using SDS-PAGE showed pfLF_highFe and pfLF_lowFe closely resembled cLF, while glycosylation analysis demonstrated very similar N-glycan compositions and site occupancy across low iron samples. The isoelectric points (pI) of pfLF_highFe and pfLF_lowFe were similar to cLF; however, varied slightly as a function of iron saturation, as did surface hydrophobicity (S_0_) measurements. Thermal denaturation analyses revealed characteristic behaviour of bovine lactoferrin preparations, with differences again observed according to the preparation’s iron saturation. Functional studies highlighted the iron chelating ability of native-LF preparations, cLF and pfLF_lowFe, and accompanying antimicrobial activity, but comparable interaction capacity with Lipid A (LPS) demonstrates preservation of structural and certain functional capabilities in precision-fermented bovine lactoferrin irrespective of iron saturation levels.

## 1 Introduction

The emergence of precision fermentation (PF) has been highlighted as a transformative technology for the food industry, with promise to revolutionise the production of complex organic materials for a myriad of use cases. Recent developments have amplified the capabilities of the technique, enabling the production of a diverse array of materials far beyond its original scope (Augustin et al., 2023). PF builds upon traditional microbial fermentation by using similar ‘microbial hosts’ such as yeast, bacteria and fungi. This process is then elevated through meticulous alterations at the genetic level to introduce specific biosynthetic capabilities. By effectively ‘reprograming’ these microorganisms, their typical metabolic pathways are overridden and refocussed on the production of specific compounds (Knychala et al., 2024). The control afforded to the producer not only unveils various possibilities from a manufacturing standpoint but may also foster a future where abundance, sustainability and productivity coalesce.

PF has been highlighted as a key pillar of the fourth industrial revolution in the food industry, offering a sustainable alternative to current unsustainable practices (Hassoun et al., 2022). By introducing substantial efficiencies into the use of land, water and feedstock resource use, PF offers a solution to growing protein demands while minimising environmental concerns like greenhouse gas emissions and soil depletion (Henchion et al., 2017; Tubb & Seba, 2019). A notable application of PF is the production of high value animal proteins such as bovine lactoferrin (bLF), a whey protein found in bovine milk and colostrum (Superti, 2020). bLF has considerable therapeutic potential, demonstrating antibacterial, antifungal, antiparasitic, anti-inflammatory and immunomodulatory properties (Yami et al., 2023).

bLF is a glycosylated protein with a mass of ∼80kDa, consisting of 689 amino acids, carrying a net-positive charge and an isoelectric point of pH 8-9 (García-Montoya et al., 2012; Jańczuk et al., 2023; Wakabayashi, Yamauchi, & Takase, 2006). It is comprised of a single polypeptide chain that is folded into two globular lobes, an N-lobe and a C-lobe, with each lobe linked by a hinge region formed by parts of an alpha helix (González-Chávez, Arévalo-Gallegos, & Rascón-Cruz, 2009). Each lobe binds one ferric ion (Fe^3+^) in coordination with a carbonate ion (CO_3_^2-^). Three classifications have been proposed based on iron saturation: apo-bLF (free of Fe^3+^), native-bLF (iron saturation levels are ∼10-30%), and fully saturated holo-bLF (Baker & Baker, 2004).

The presence or absence of bound iron plays a critical role in determining the structural orientation and functionality of bLF. Key characteristics such as isoelectric point (pI), thermal stability and tertiary structure are known to vary as a function of iron saturation (Bokkhim et al., 2013; Wang et al., 2013). Lactoferrin’s iron chelating ability also limits the amount of free iron in bodily fluids and tissues, therefore inhibiting the growth of pathogens that require iron (Rosa et al., 2017). This capability becomes diminished as the protein reaches elevated levels of iron saturation, underscoring the functional differences that may manifest according to the state of bLF. Iron chelation, however, is not the only feature of bLF that drives its bio-functionality. It is now well understood that lactoferrin’s structure enables direct interactions with various strains of bacteria, destabilising their membranes and facilitating their breakdown (Ling & Schryvers, 2006). bLF has also been shown to modulate the immune system by influencing cytokine production and enhancing the activity of immune cells, contributing to its broad-spectrum antipathogenic activity (Legrand et al., 2005).

More recent findings also highlight the importance of post-translational modifications (PTMs) such as glycosylation in maintaining structural integrity and facilitating protein function (Adlerova, Bartoskova & Faldyna, 2008). The best characterised form of glycosylation in bLF is N-linked glycosylation, whereby sugar units bind at the nitrogen atom of asparagine (Asn) residues. Five possible sites of attachment include Asn 233, 281, 368, 476 and 545, however only 15-30% of bLF molecules show this glycan attachment at Asn 281 (Karav et al., 2017). A complete understanding of the relationship between N-glycosylation and protein structure and function in bLF is still lacking, however there does seem to be some critical interplay. Legrand et al. (1990) demonstrated a large reduction in the iron binding capacity of deglycosylated bLF tryptic fragments, suggesting a link between glycosylation and iron chelation, and therefore the protein’s bio-functionality. van Veen et al. (2004) also found that bLF molecules with attachments at Asn281 had an increased proteolytic resistance, afforded by steric hinderances that restrict access of proteases to Lys282. Replication of these glycan structures in PF bLF therefore appears critical to preserve the broad functional capabilities of the naturally occurring protein.

While the synthesis of bLF through PF heralds a new frontier in sustainable protein production, it introduces a layer of complexity: although the technique ensures production of the target protein at a primary structure level, variability in host-cell machinery may result in divergences in higher-order structures (Chen, 2011). Given that deviations from structural fidelity may manifest as functional differences, rigorous physicochemical and functional comparisons are necessary. We therefore aim to provide a structural and functional comparison between a commercial bLF (cLF) and two precision-fermented bLF samples of varying iron saturation levels, pfLF_highFe and pfLF_lowFe, produced from the same *Pichia pastoris* production strain. Properties such as molecular weight, pI, molar absorptivity, glycosylation and surface hydrophobicity will be evaluated. These comparisons will be integrated with functional analyses of antibacterial and anti-inflammatory properties, to holistically assess the efficacy of the precision fermentation process. Such characterisation not only establishes bioequivalence, but also predicates its efficacy as a bioactive nutraceutical for commercial use.

## 2 Materials and Methods

### 2.1 Materials

Tris-base and Tris-HCl powders (purity >99.0%) were sourced from Sigma-Aldrich (St. Louis, MO) and dissolved in Milli-Q water (Millipore Corporation, VT) to prepare a Tris buffer solution. bLF for amino acid sequencing was purchased from Sigma-Aldrich (St. Louis, MO), food-grade, commercial spray dried bovine lactoferrin (cLF) powder was purchased from the Lactoferrin Co. (batch # 21240G, 95% lactoferrin) and provided by All G (Sydney, Australia), along with two precision-fermented variants (pfLF_highFe and pfLF_lowFe) differing only in iron saturation levels. 8-anilino-1-naphthalenesulfonic acid (ANS), *E. Coli,* Lipid A natural bisphosphoryl, goat anti-rabbit IgG-Peroxidase, and SIGMA*FAST*™ OPD were from Sigma-Aldrich (St. Louis, MO), while anti-LF was from Thermo Fisher (Waltham, MA). All other chemicals were of analytical grade, and solutions were prepared and diluted in Tris-buffer on the day of experimentation. Experiments were completed in triplicate unless specified otherwise.

### 2.2 Methods

#### 2.2.1 Precision Fermentation of bovine lactoferrin

Precision fermented lactoferrin (pfLF) is produced through fermentation of a strain of *Komagataella phaffii* (originally called *Pichia pastoris*) that has been modified to secrete recombinant bovine lactoferrin into the fermentation media*. K. phaffii* BG11 was used to develop the pfLF production strain and is derived from a lineage of *K. phaffii* (NRRL Y-11430) that is well-characterised, and has a long history of safe use in the manufacture of various food enzymes and pharmaceutical agents (Cereghino and Cregg, 2000). Modifications to *K. phaffii* BG11 were introduced to increase the titre of pfLF and modifications of the glycosylation pathway were also introduced to ensure the pfLF also contains the main glycans found in native bLF. The fermentation conditions of the strain are confidential, but to control the iron saturation of the produced pfLF, specific concentrations of FeSO4 in the fermentation medium were used. After fermentation, cells were separated from the pfLF containing liquid before the pfLF was purified by cation-exchange chromatography. The eluted pfLF was then concentrated and filtered to remove the salt from elution before being freeze dried for analysis.

#### 2.2.2 SDS-PAGE

Solutions of lactoferrin preparations were prepared by mixing 10 µL of sample with 10 µL of BioRad 2x Laemmli sample buffer containing 100 mM DTT, to a final concentration of 50mM DTT. Mixtures were heated at 95°C for 5 min before 10 µL of sample solution was added to wells. Proteins were resolved on Mini-PROTEAN TGX Stain-Free Gels (7.5–20%) in Tris/Glycine/SDS buffer at 180 V for 30 min. 4 µL of Precision Plus Protein Unstained Standards were loaded as markers. Gels were imaged using a Bio-Rad Gel Doc imager with the “Stain-Free Gel” setting.

#### 2.2.3 Purity using Independent Data Acquisition (IDA)

For bottom-up experiments, lactoferrin powder samples were diluted in 25 mM ammonium bicarbonate (NHLHCOL), and protein concentration was measured using a NanoDrop spectrophotometer. 50 μg of each sample was reduced with 2 mM DTT for 30 minutes, followed by protein alkylation using 11 mM chloroacetamide (CAA) and incubation in darkness for 30 minutes. Trypsin, at a 1:50 (enzyme:substrate) ratio was added, and samples incubated for 18 h at 37°C. The enzymatic reaction was stopped with trifluoroacetic acid (TFA) at a final concentration of 0.2%. 1 μg of digested peptides was injected into a PROTECOL C18 G 203 (250 mm x 300 μm) analytical column for LC-MS analysis.

Purity was analysed in Maxquant v2.4.7.0, with search parameters including carbamidomethylation of cysteines (fixed), oxidation of methionine, and N-terminal acetylation (variable). Analysis parameters included trypsin, chymotrypsin or none as the digestive enzyme, a peptide tolerance of 10 ppm, a false discovery rate (FDR) of 0.01, two missed cleavages, a peptide length of 7–45 amino acids, and site decoy fraction of 0.01. To be considered a contaminant, proteins must be identified and quantified in at least two of three replicates, with quantification based on the average percentage.

#### 2.2.4 Amino acid sequence coverage and glycosylation site occupancy calculation

Lactoferrin samples were diluted in 25 mM ammonium bicarbonate (NHLHCOL), and protein concentration was measured at 280 nm. Each 25 μg sample was reduced with DTT (final concentration 2 mM) for 30 minutes, followed by alkylation with CAA (final concentration 11 mM) in darkness for 30 minutes. Trypsin, elastase, and chymotrypsin were added separately (1:50 enzyme:protein), and samples incubated for 18 h at 37 °C. Reactions were stopped with TFA (final concentration 0.2%). Digested peptides were analysed by LC-MS/MS on a ZenoTOF™ 7600 with a Waters Microscale LC System. Peptides were separated chromatographically using a 25-minute microflow gradient (5–60% phase B) at 7 µL/min with mobile phases of 0.1% v/v formic acid in water (A) and 0.1% v/v formic acid in acetonitrile (B).

ProteinPilot™ Software 5.0.2 was used to analyse amino acid sequence coverage while PEAKS Studio 12 for glycosylation site occupancy. Parameters included 10 ppm parent mass error, 0.02 Da fragment mass error, and monoisotopic precursor mass search. Enzyme specificity, semi-specific digestion and peptide lengths of 6–45 amino acids were set. For amino acid sequencing, Sigma bLF was used arather than cLF as a comparative baseline since it provides a well characterised standard. Fixed modifications were set to carbamidomethylation (+57.02 Da), and variable modifications included oxidation of methionine (+15.99 Da). A maximum of two PTMs per peptide was allowed. Deep learning-based FDR estimation was enabled with a confidence amino acid threshold of 2.00%. N-glycan composition and glycosite occupancy was thus completed.

#### 2.2.5 Free glycan analysis

Proteins were dissolved in water and diluted to 10 mg/mL. Aliquots of 20 µg were reduced, alkylated, spotted onto PVDF membranes and air-dried. Membranes were blocked with 1% (w/v) polyvinylpyrrolidone (PVP) and washed. N-glycans were released using 10 U PNGase F (Promega) at 37 °C overnight. N-glycans were deaminated, dried, reduced with sodium borohydride and cleaned with graphite carbon-packed tips. O-glycans were released by reductive beta-elimination and similarly cleaned. A total of 10 µg glycan equivalents were injected into the LC-MS, analysed in negative ion mode (m/z 400–2000) and the top five most intense ions were fragmented. Glycans were separated using a Hypercarb column (1 mm ID × 30 mm, 3 µm particle size; ThermoFisher) with a gradient of Buffer A (10 mM ammonium bicarbonate in water) and Buffer B (10 mM ammonium bicarbonate in 70% ACN). Gradient elution information is available in **Supplementary Table 1.**

#### 2.2.6 Iron saturation analysis by UV-vis

Iron saturation was compared across lactoferrin preparations using UV-vis spectroscopy. The UV-vis method for iron saturation relies on the ratio of absorbance of lactoferrin at 280 nm and 465 nm using E280 = 1.51 and E465 = 0.57 (1% and 100% iron saturation, respectively) (Foley & Bates, 1987; Groves, 1960; Majka et al., 2013). Lactoferrin powders were dissolved in 5 mM Tris-HCl with 150 mM NaCl at 10 mg/mL, then diluted to 1 mg/mL. Measurements (600–260 nm) were taken using a Shimadzu 1900i UV-vis spectrophotometer with a TMSPC-8 cell holder, 1 cm pathlength and 150 μL sample volume. Iron saturation was calculated from absorbance ratios of E280/1.51 and E465/0.57.

#### 2.2.7 Iron binding capacity

The iron binding capacity of lactoferrin samples was examined following the method of Chen and Wang (1991). Iron binding capacity was determined by taking the ratio of A465/0.57 at 0.3 mM Fe^3+^. A Shimadzu 1900i UV-vis spectrophotometer equipped with a TMSPC-8 cell holder for an 8 cell microcuvette with a 1 cm pathlength and 150 μL sample volume was used for measurements between 600-260 nm.

#### 2.2.8 UV-vis spectroscopy

The absorbance profiles and extinction coefficients (ε) of lactoferrin preparations were examined using UV-vis spectroscopy. Solutions were prepared in 10 mM Tris buffer (pH 7.2), with concentrations systematically being varied from 0.1–0.5 mg/mL to create a calibration curve. Measurements at 280 nm were performed using a Lambda 35 spectrophotometer (Perkin Elmer, Washington, United States) and a 1 cm path length quartz cuvette. Absorbance values were plotted against concentrations, and the ε value was calculated from the gradient of the linear relationship.

#### 2.2.9 Fluorescence spectroscopy

Fluorescence emission spectra for pH 7.2 lactoferrin samples were captured using a Fluromax-4 Spectrophotometer (Horiba Scientific, Kyoto, Japan). Solutions were analysed using a 1 cm quartz cuvette at a concentration of 0.5 mg/mL. Measurements were made at 25°C, with an excitation wavelength of 295 nm. Emission spectra were acquired over a spectral range of 305 to 400 nm, with emission slit widths being set to 3 nm.

#### 2.2.10 Isoelectric point (pI)

The zeta potentials of the lactoferrin preparations were determined using a Zetasizer (Malvern Instruments Ltd, Malvern, UK). 0.5 mg/mL solutions of both proteins were prepared in Milli-Q water, and the pH of the preparations was adjusted using small aliquots of 0.1 M NaOH. The refractive index used for the protein preparation was 1.50 and 1.33 for the dispersant (Milli-Q water).

#### 2.2.11 Fourier transform infrared spectroscopy (FTIR)

FTIR spectra of pH 7.2 lactoferrin samples were acquired using a Spectrum 2 FTIR spectrometer (Perkin Elmer, Norwalk, CT, United States) at a concentration of 0.5 mg/mL. Analysis employed a GladiATR (Pike Technologies, Madison, WI, United States) diamond crystal attenuated total reflectance (ATR) device. Samples were analysed over a narrowed spectral region of 4000–400 cm^−1^ under a constant nitrogen purge. Spectra represented an average of 100 scans at a resolution of 4 cm^−1^, executed at 25°C. Tris-buffer artifacts were subtracted from interferograms using the least squares method. Secondary structure contributions were estimated using OPUS 8.2 software (Bruker Corporation, Billerica, United States), as previously detailed by Condict et al., 2019.

#### 2.2.12 Circular dichroism

Lactoferrin samples were analysed at a concentration of 0.5 mg/mL and a pH of 7.2. The CD measurements were executed using a Jasco (J-1500) spectropolarimeter (Jasco International Co. Ltd., Tokyo, Japan), at ambient temperature and under a continuous nitrogen flow. Spectral data were collected in the far UV-region, spanning from 190 to 260 nm, using a cylindrical quartz cell with an optical path length of 0.1 mm. Three accumulations were taken under a scan rate of 50 nm/min, a data pitch of 1 nm, a bandwidth of 1.0 nm and averaged. Spectra were then analysed using the CONTINN method, and data set 4 was used for estimation of the secondary structure *via* the DichroWeb platform (http://dichroweb.cryst.bbk.ac.uk).

#### 2.2.13 Surface hydrophobicity

Investigation into the surface hydrophobicity of protein preparations was conducted employing an adapted ANS fluorescent probe method as described by Alizadeh-Pasdar & Li-Chan (2000). Serial dilutions of protein from 0.005% to 0.03% (w/v) were prepared for cLF and pfLF_lowFe, and from 0.03% to 0.1% (w/v) for pfLF_highFe to facilitate protein-ANS complexation. 20µL aliquots of ANS were added to each dilution, alongside a blank protein sample of the same concentration. The relative fluorescence intensity (RFI) was measured for each dilution with excitation and emission wavelengths of 390 nm and 470 nm, respectively.

#### 2.2.14 Thermal stability via differential scanning calorimetry

The thermal stability and denaturation enthalpies of all lactoferrin preparations were determined using a differential scanning calorimeter Q2000 (TA Instruments, New Castle, DE, USA). Samples were prepared at 10 mg/mL in Tris Buffer (pH 7.2), and 5 μL of solution was hermetically sealed in Tzero pans. Preparations were equilibrated at 10°C for 30 minutes, then analysed at a scan rate of 3°C/min, from 10°C–120°C under nitrogen flow. From DSC transition peak data, thermodynamic parameters were determined, including the maximum heat absorption temperature (T_max_), onset temperature (T_s_), and denaturation enthalpy change (ΔH_cal_).

#### 2.2.15 Antimicrobial assay

To test the antimicrobial properties of lactoferrin against E. coli, a modified method based on Sekse et al. (2012) was used. *E. coli* strain W3110 (K12) was tested for growth inhibition with lactoferrin concentrations from 0.0125 to 4 mg/mL. 2X 5.25 g/L Mueller-Hinton (MH) broth was prepared in MilliQ water and autoclaved. Precultures were grown overnight at 37°C, diluted into fresh MH to an OD of 0.025, and 200 μL inoculated into a sterile 2 mL 96 deepwell plate. Lactoferrin solutions (20 mg/mL in MilliQ water) were adjusted to 4 mg/mL, and a dilution series was prepared. 200 μL of each lactoferrin concentration was added to 200 μL cultures and incubated for 16 h at 37°C with shaking (1000 RPM, 3 mm throw). 100 μL samples were transferred to a Greiner cat. # 655061 96-well plate, and absorbance at 600 nm was measured using a Byonoy Absorbance 96 Automate plate reader. Appropriate blanks were subtracted.

#### 2.2.16 ELISA assay

An enzyme-linked immunosorbent assay (ELISA) was employed as described by Appelmelk et al. (1994) with some modifications, including the use sodium azide to replace merthiolate. In brief, plates were first coated with *E. coli*, lipid A natural bisphosphoryl, followed by lactoferrin at varying concentrations, prior to the addition of anti-LF and finally anti-rabbit IgG-peroxidase conjugated. Following incubation and deactivation of the enzyme with 10% sulfuric acid, absorbance was measured at 492 nm using a FluoSTAR Omega microplate reader (BMG Labtech, Germany). A control containing only lactoferrin was included to ensure binding to plastic did not influence results.

#### 2.2.17 Statistical Analysis

Analyses were performed in triplicate and results were expressed as mean ± standard error. Data was analysed by one-way ANOVA with post-hoc Tukey HSD (Honestly Significant Difference) performed with Graph pad Prism 9.4.1 software (GraphPad software, USA), with a p value of ≤ 0.05 being considered statistically significant.

## 3 Results and discussion

### 3.1 Primary Structure and sequence coverage

The pfLF gene was designed from the mature protein sequence without the signal peptide of bLF from the UniProt database (UniProt P24627) and codon optimised for *P. pastoris.* To confirm the correct protein identity of pfLF preparations, bottom up LC-MS/MS utilised three different proteases, trypsin, elastase, and chymotrypsin to ensure full amino acid sequence coverage. pfLF_lowFe and pfLF_highFe were confirmed to have 100 % amino acid sequence coverage compared to the native Sigma bLF with 917, 661 and 864 unique peptides analysed, respectively (**Supplementary Table 2,3,4,5**).

### 3.2 SDS-PAGE (intact) - purity, glycosylation analysis

SDS-PAGE (**Figure 1**) was used to compare the electrophoretic migration of pfLF_lowFe and pfLF_highFe with cLF and showed that the proteins migrate to the same molecular weight of ∼75kDa. The purity of the LF samples was investigated by IDA LC-MS/MS with pfLF_lowFe, pfLF_highFe and cLF being 99 %, 98.6 % and 99 % pure, respectively.

**Fig. 1.**
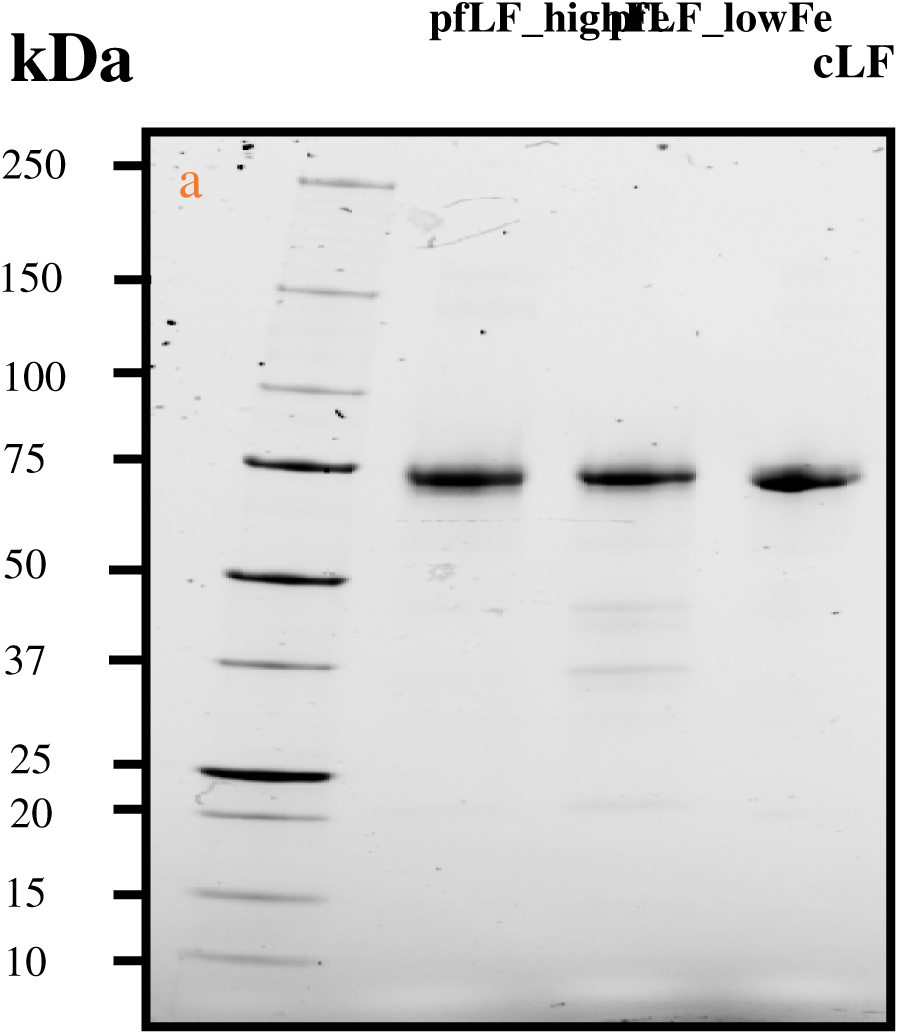
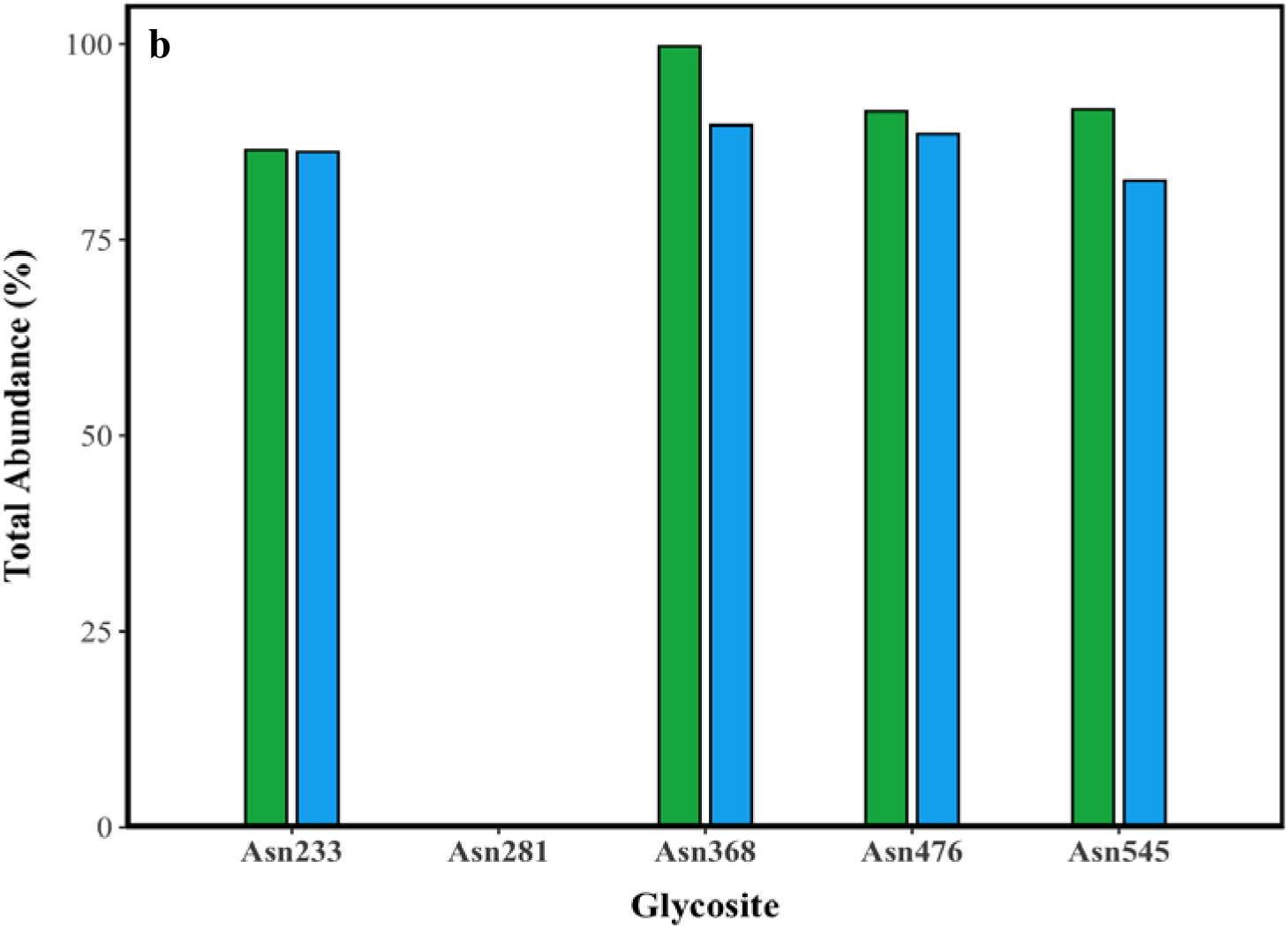
(a) SDS-PAGE analysis of intact cLF, pfLF_highFe and pfLF_lowFe. (b) Relative abundance (%) of glycosite occupancy at positions Asn233, Asn281, Asn368, Asn 476 and Asn545 for cLF (green) and pfLF_lowFe (blue).

Glycosite occupancy was also calculated using IDA LC-MS/MS. Only pfLF_lowFe glycans were analysed as this is the protein being commercialised, and both pfLF proteins originate from the same strain. Results indicate that pfLF_lowFe is glycosylated the same or slightly less than cLF at positions Asn233, Asn368, Asn476 and Asn545 (**Figure 1b**). Glycans were not identified at Asn281 in either pfLF_lowFe or cLF.

LC-MS/MS was also used to analyse the type and total abundance of glycans that were identified for pfLF_lowFe and indicate that the attached glycans have a range of 7 to 13 mannose residues, with the highest abundances being 8 (33.38 %) and 9 (32.35 %) mannoses, consistent with the highest abundances observed for cLF (**Table 1**) and also with literature values (Valk-Weeber et al., 2020). In addition, 23.73% of the total glycans contained 1 phosphate group, although only in the range of 9-13 mannose residues was observed to be phosphorylated. This amount of phosphomannose was the same as that previously observed for recombinant human lactoferrin produced by *K. phaffii* (Lu et al., 2024) and consistent with the amount of phosphomannose observed in other native and recombinant proteins produced by K. phaffii (Grinna and Tschopp, 1989; Miele et al., 1997; Montesino et al., 1998; Miura et al., 2004).

**Table 1.**
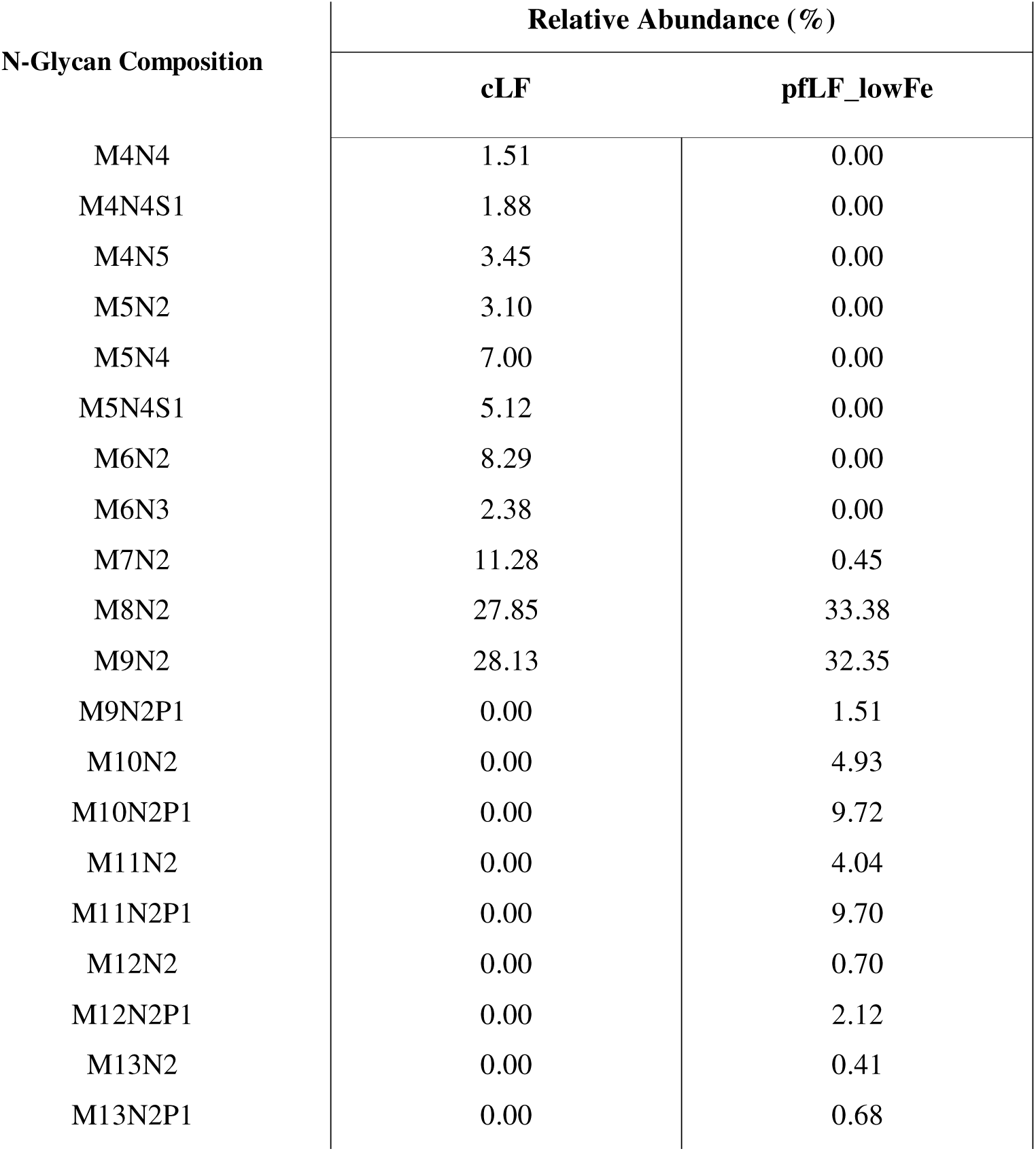
Relative abundance (%) of N-glycan compositions in cLF and pfLF_lowFe. N-glycan structures are described using shorthand notations, where “N” represents Mannose-N-Acetylated, “M” indicates Mannose, “P” denotes Phosphate, and “S” refers to N-Acetyl Neuraminic Acid.

| N-Glycan Composition | Relative Abundance (%) |  |
| --- | --- | --- |
|  | cLF | pLF_lowFe |
| M4N4 | 1.51 | 0.00 |
| M4N4S1 | 1.88 | 0.00 |
| M4N5 | 3.45 | 0.00 |
| M5N2 | 3.10 | 0.00 |
| M5N4 | 7.00 | 0.00 |
| M5N4S1 | 5.12 | 0.00 |
| M6N2 | 8.29 | 0.00 |
| M6N3 | 2.38 | 0.00 |
| M7N2 | 11.28 | 0.45 |
| M8N2 | 27.85 | 33.38 |
| M9N2 | 28.13 | 32.35 |
| M9N2P1 | 0.00 | 1.51 |
| M10N2 | 0.00 | 4.93 |
| M10N2P1 | 0.00 | 9.72 |
| M11N2 | 0.00 | 4.04 |
| M11N2P1 | 0.00 | 9.70 |
| M12N2 | 0.00 | 0.70 |
| M12N2P1 | 0.00 | 2.12 |
| M13N2 | 0.00 | 0.41 |
| M13N2P1 | 0.00 | 0.68 |

### 3.3 Iron Saturation/Iron Content

The iron saturation of cLF, pfLF_highFe and pfLF_lowFe was measured using UV-vis and shown to be 18.3 %, 96.3 % and 15.8 %, respectively. The iron binding capacity of cLF, pfLF_highFe and pfLF_lowFe was then also measured using UV-vis spectroscopy (**Figure 2**). Iron binding capacities were calculated to be 65.70 % (cLF), 10.67 % (pfLF_highFe) and 77.65 % (pfLF_lowFe), in line with the initial iron saturation status. The ability to manipulate the iron saturation and hence iron binding capacity of pfLF by altering the amount of FeSO_4_ in the fermentation medium towards a desired protein functionality, e.g. a source of bioavailable Fe^3+^ (high saturation) (Zhao et al. 2022) or the ability to sequester Fe^3+^ for bioactivity such as antimicrobial activity (low saturation), shows the flexibility of precision fermentation to produce tailored protein products depending on the end-use.

**Fig. 2.**
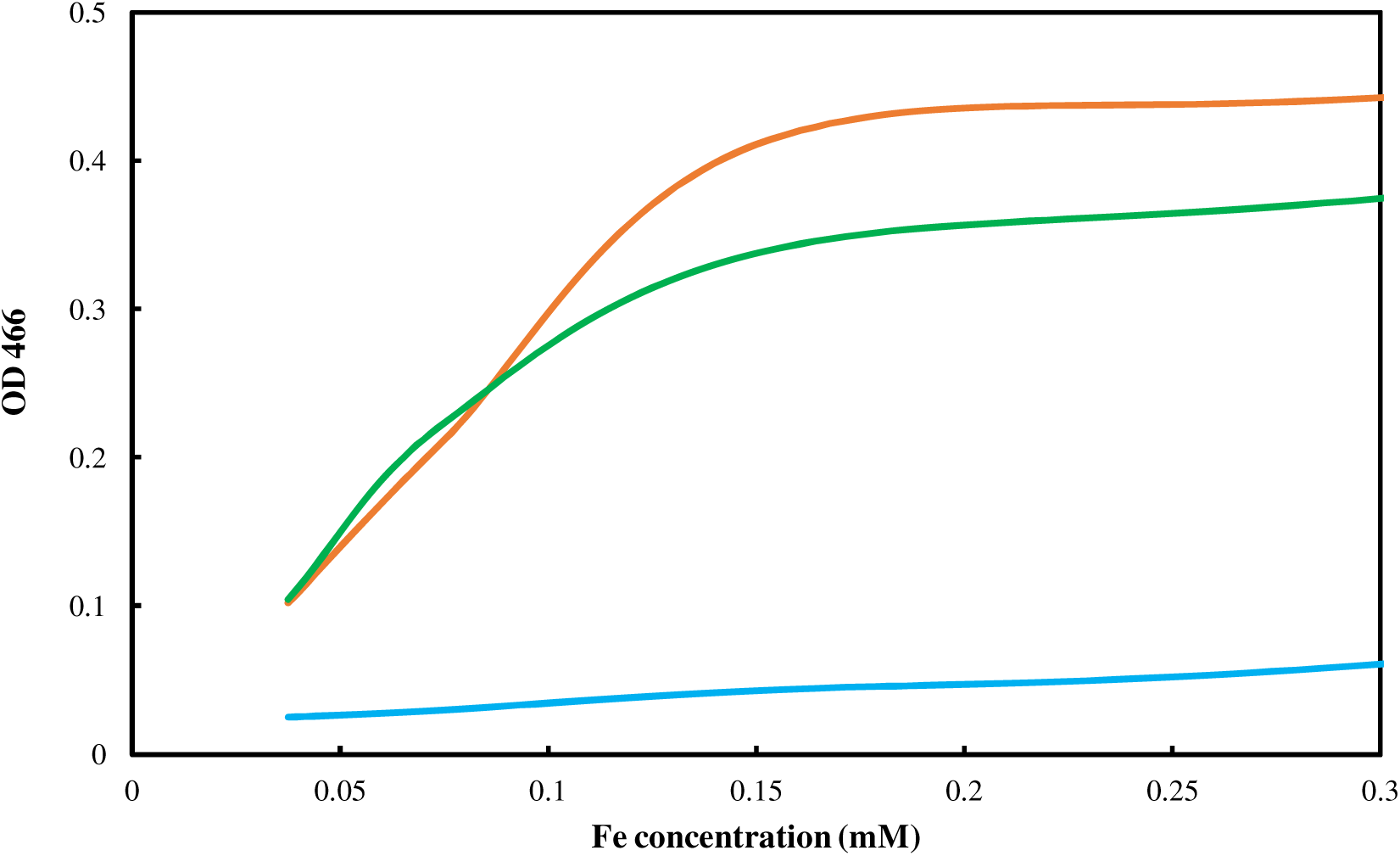
UV-vis absorbance at 466 nm of 10 mg/mL cLF (green), pfLF_highFe (orange) and pfLF_lowFe (blue) at varying Fe^3+^ concentrations. The ratio of measured A_466/_0.57 was used to determine iron binding capacity at 0.3mM Fe^3+^ where there is a slight excess of Fe^3+^.

### 3.4 Spectroscopic Analysis: Molar Extinction and Fluorescence Characterisation

The extinction coefficient, or molar absorption coefficient (ε) relates to absorbance and attenuation strength of a chemical species at a given wavelength (Aitken & Learmonth, 2002). Given this value is an intrinsic property of the chemical species being analysed, it allows for protein concentration to be accurately determined in downstream applications. Hence, ε at 280 nm was determined for cLF, pfLF_highFe and pfLF_lowFe by plotting absorbance at this wavelength as a function of increasing concentration (**Figure 3a**). The ε values of native-LF preparations, cLF and pfLF_lowFe, were found to be 10.16 x 10^4^ and 10.44 x 10^4^ M^-1^ cm^-1^ respectively, whereas pfLF_highFe, which has a substantially higher iron saturation, showed an ε value of 10.48 x 10^4^ M^-1^ cm^-1^. While all samples exhibited very similar ε values, the minor observed differences seem to indicate an increased molar absorptivity with higher iron saturation. The obtained values are also commensurate with previously reported values for bLF (Shimazaki et al., 1998). The very similar molar absorptivity values of the three samples suggests an identical quantity and distribution of absorbing residues across all preparations, again highlighting the efficacy of the precision fermentation process in structural conservation.

**Fig. 3.**
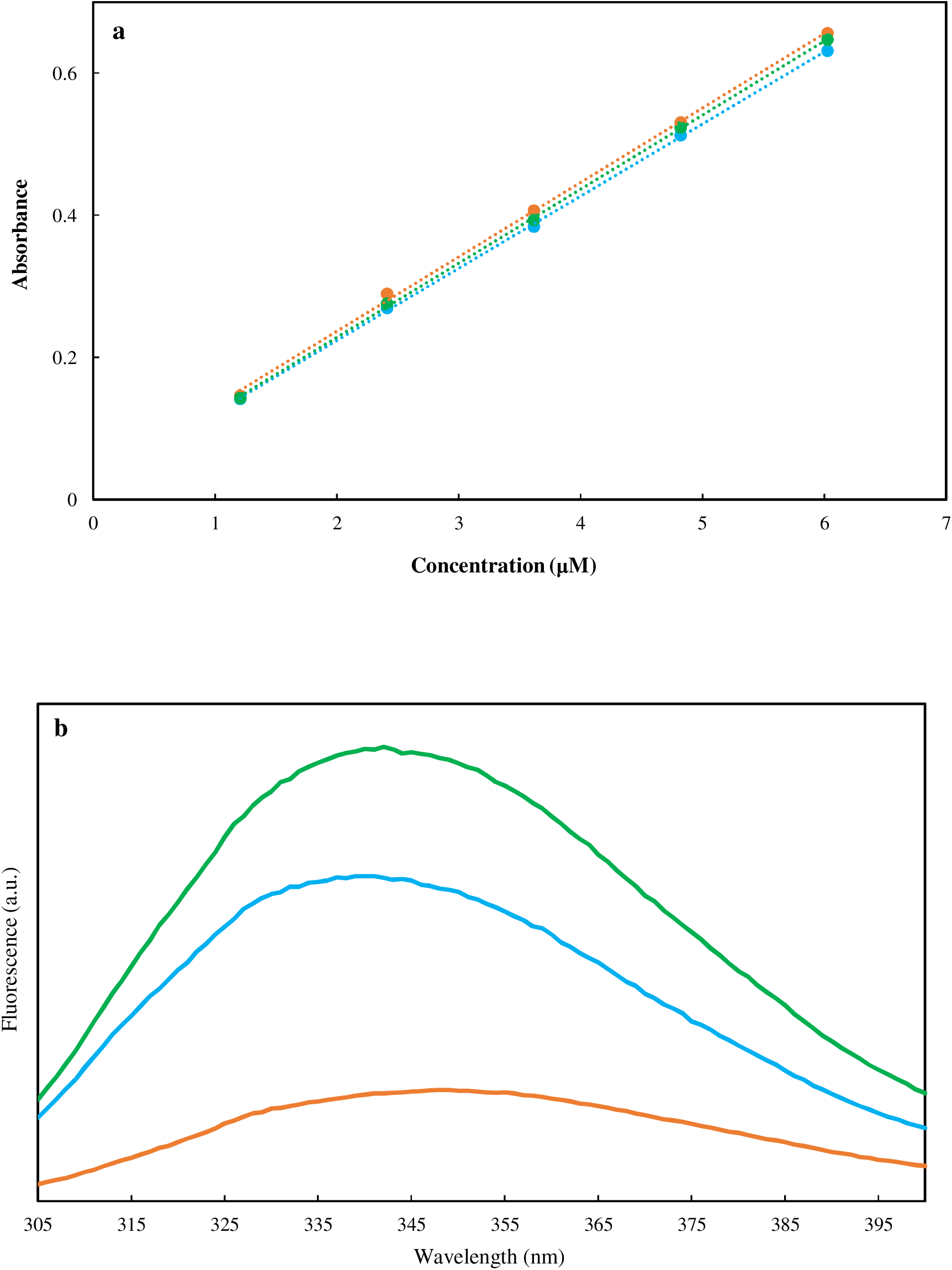
(a) UV-vis absorbance at 280 nm of cLF (green), pfLF_highFe (orange) and pfLF_lowFe (blue) across a concentration range of 0.1 mg/mL – 1 mg/mL. (b) Fluorescence spectra of 0.5mg/mL cLF (green), pfLF_highFe (orange) and pfLF_lowFe (blue) with an excitation wavelength of 295 nm and an emission wavelength ranging from 305-400 nm.

Fluorescence spectroscopy was also employed to provide additional insights on the tertiary structure of each preparation. An excitation wavelength of 295 nm was selected to exclusively excite tryptophan residues, further probing structural fidelity across preparations and the impact of iron binding on tryptophan emission. **Figure 3b** shows the fluorescence spectra of all lactoferrin preparations across an emission wavelength of 305-400 nm. When comparing the PF samples, differences in emission peaks were observed, which appear to be related to the protein’s iron saturation levels. The emission maxima were 341 and 348 nm for pfLF_lowFe and pfLF_highFe, respectively, with the observed fluorescence also being significantly reduced in the high iron sample. These observations are in close agreement with previously reported effects of iron binding on bLF fluorescence (Bokkhim et al., 2013; Wang et al., 2013).

Upon iron binding, lactoferrin is known to undergo a conformational change towards a more ‘compact’ structure. This makes an observed red shift (usually indicative of a more hydrophilic microenvironment surrounding tryptophan residues) seem counterintuitive. However, the absorbance of emission light by the Fe^3+^ molecules is likely the major contributor to this apparent red shift, rather than a change induced by a conformational alteration. Similar effects have been shown in iron-transferrin systems by Lehrer (1969) who proposed a Förster Resonance Energy Transfer (FRET) as the mechanism behind the observed quenching to the red side of the emission spectrum, arising from the spectral overlap between donor (tryptophan) emissions and acceptor (Fe-transferrin) absorption bands.

Despite a similar iron saturation to pfLF_lowFe, cLF exhibits the greatest fluorescence intensity. This is likely due to differences in drying methods, with the cLF sample appearing to be spray-dried, while precision-fermented samples were lyophilised. Hence, the thermal treatment applied to cLF may be responsible for differences in intrinsic fluorescence. Stănciuc et al. (2013) also note an increase in bLF intrinsic tryptophan fluorescence upon thermal denaturation (80°C for 10 minutes). In the native state, the thirteen bLF tryptophan residues are expected to experience intramolecular quenching by surrounding aromatic residues, however upon thermal denaturation, distances between tryptophan residues and intramolecular quenchers are significantly altered, thus impacting the observed intrinsic fluorescence intensity of the protein. This possibility, along with implications for thermal stability and surface hydrophobicity, is explored further in subsequent sections.

### 3.5 Isoelectric Point

The isoelectric point (pI) is a key comparative parameter between cLF and its precision-fermented counterparts, reflecting the pH at which a protein carries no net charge. This affects solubility, formulation compatibility and potential therapeutic applications. Zeta-potential analysis identified the pH at which each preparation exhibited net zero charge, indicative of their pI. **Figure 4a-c** shows zeta-potential as a function of pH for cLF (**Figure 4a**), pfLF_highFe (**Figure 4b**), and pfLF_lowFe (**Figure 4c**). Initial pH values in Milli-Q water were 5.77, 6.32 and 6.30, respectively, demonstrating a predominantly positive charge which is consistent with existing reports (Baker & Baker, 2004).

**Fig. 4.**
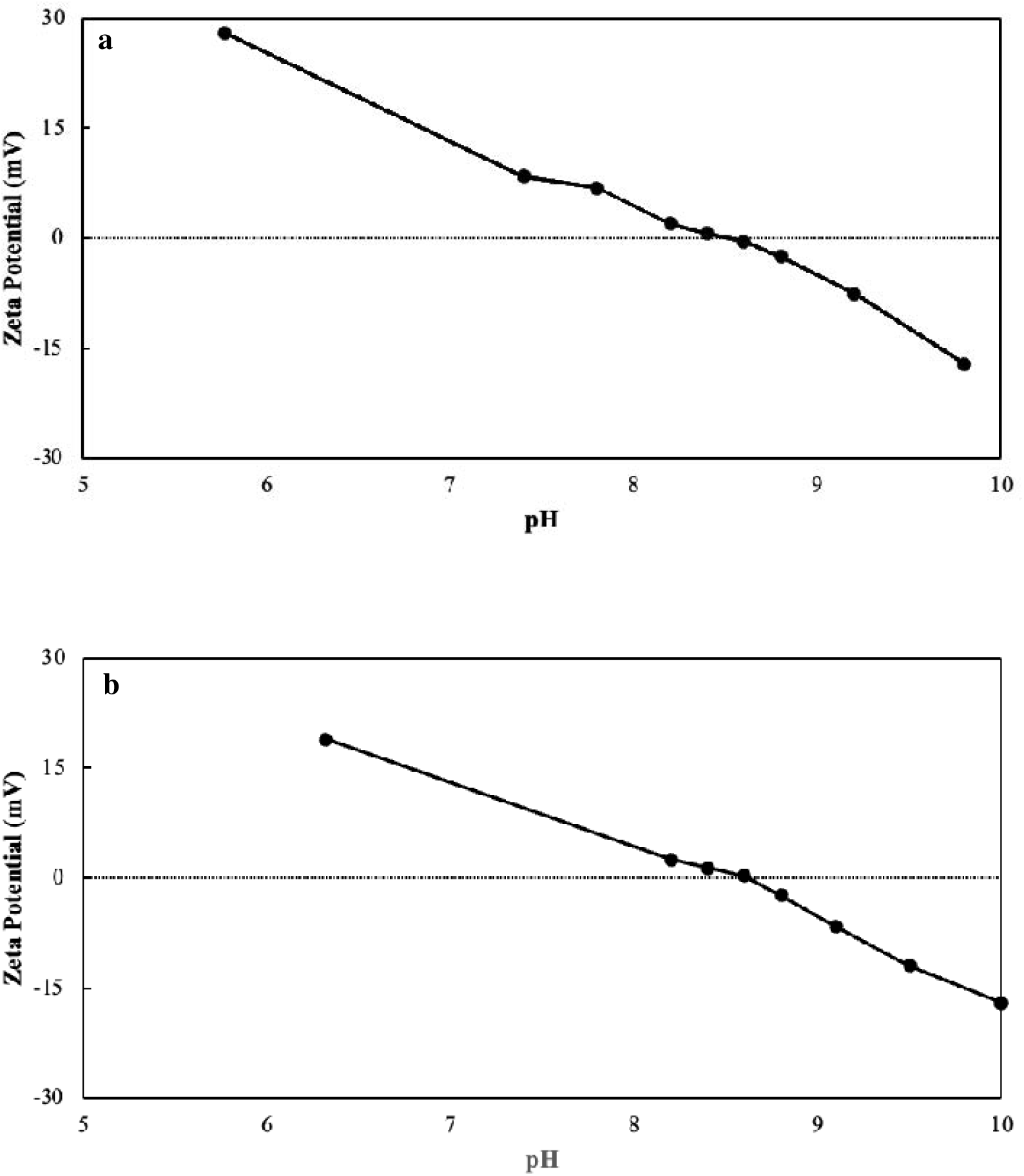
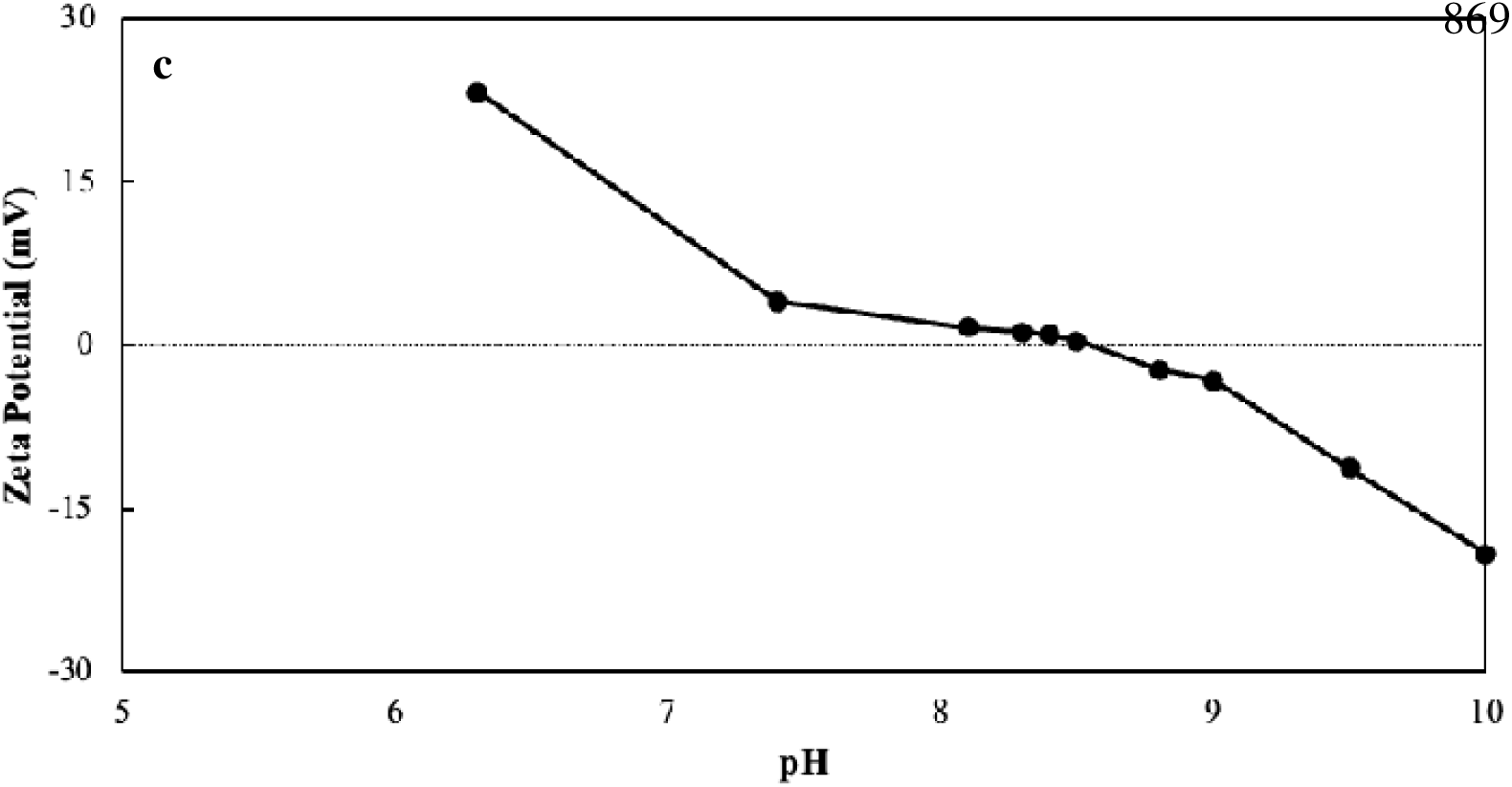
Zeta potential (mV) of 0.5 mg/mL cLF (a), pfLF_highFe (b) and pfLF_lowFe (c) as a function of increasing pH.

It also appears that iron saturation impacts surface charge. Native-LF preparations (cLF and pfLF_lowFe) with iron saturation of 15.79% and 18.29% respectively, showed ζ approaching 0 mV at ∼pH 8.5, while holo-LF pfLF_highFe with 96.26% iron saturation reached 0 mV at ∼pH 8.7. These measurements are consistent with reported literature values for the pI of bLF, which range between 8 and 9 (Dyrda-Terniuk & Pomastowski, 2023) and reflect conformational differences due to iron binding. Iron-bound lactoferrin likely buries ionisable amino acids as it adopts a more compact orientation, resulting in a higher pI. Meanwhile, native-LF may show increased exposure of these residues, lowering the pI. Additionally, glycan structures and chemically charged groups at glycan termini influence recorded pI values, contributing to observed differences between samples (Barrabés et al., 2010).

### 3.6 Secondary structure estimations

Conservation of secondary structure is also critical in maintaining protein stability and function. Precise alignment in structures such as alpha-helices, beta-sheets and beta-turns ensures that the biological activities and interaction capabilities of the protein are preserved (Rehman, Farooq & Botelho, 2023). To examine potential differences in the secondary structure of cLF and the precision-fermented samples, alterations in the amide bands were investigated. Specifically, the amide I region (1700–1600 cm^−1^), which primarily reflects the C=O stretching vibration of peptide linkages, and the amide II region (1600–1500 cm^−1^), arising from C-N stretching and N-H bending vibrations, were analysed to estimate relative structural contributions.

In the 1700–1500 cm^−1^ spectral region of all lactoferrin preparations, two significant peaks corresponding to the amide I and II regions of the protein were identified (**Figure 5a**), with the former being the primary focus of the investigation. Each of the samples displayed very similar amide I absorbances, hence, quantitative analysis was performed *via* second derivative resolution enhancement followed by curve fitting to reveal the contribution of overlapping absorption bands, characteristic of different secondary structure elements. Results of this analysis are presented in **Table 2**.

**Fig. 5.**
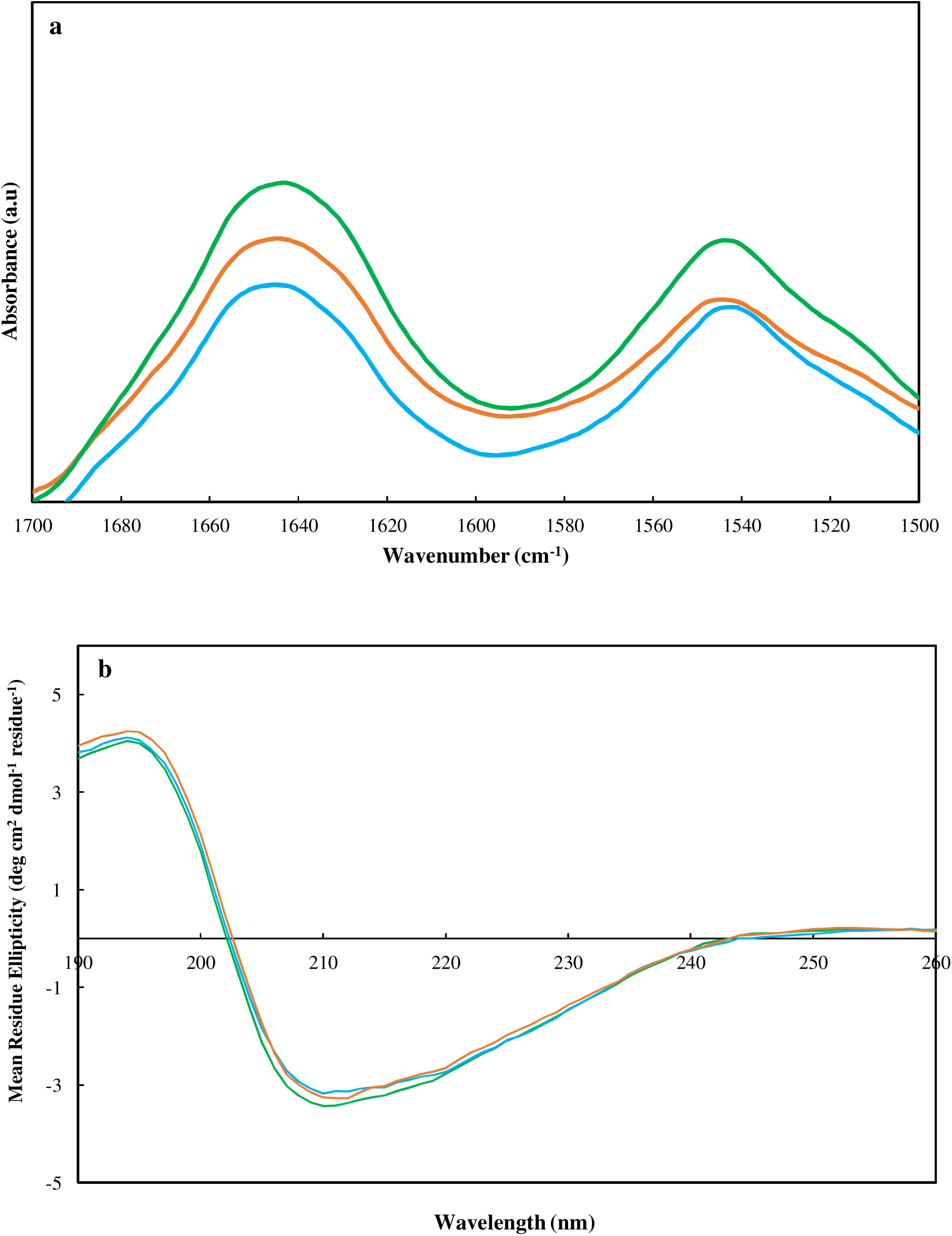
(a) FTIR spectra in the 1500-1700 cm^-1^ region (amide I and amide II) for 0.5 mg/mL cLF (green), pfLF_highFe (orange) and pfLF_lowFe (blue). (b) Circular dichroism (CD) spectra for 0.5 mg/mL cLF (green), pfLF_highFe (orange) and pfLF_lowFe (blue) across wavelengths of 190-260 nm.

**Table 2.** Fourier-Transform Infrared Spectroscopy (FTIR) and Circular Dichroism (CD) secondary structure component estimations of cLF, pfLF_highFe and pfLF_lowFe at pH 7.2.

| Secondary structure component | cLF | pfLF_highFe | pfLF_lowFe |
| --- | --- | --- | --- |
| **FTIR** |  |  |  |
| **α-helix (%)** | $20.30 \pm 0.20^a$ | $18.78 \pm 0.59^{a,b,d}$ | $20.50 \pm 0.45^a$ |
| **β-sheet (%)** | $29.50 \pm 0.26$ | $32.27 \pm 1.10^d$ | $30.11 \pm 0.92^{b,c}$ |
| **β-turns (%)** | $13.50 \pm 0.10^a$ | $11.78 \pm 0.51^a$ | $12.83 \pm 0.40$ |
| **Unordered (%)** | $36.70 \pm 0.56$ | $37.17 \pm 1.24$ | $36.56 \pm 0.64$ |
| **CD** |  |  |  |
| **α-helix (%)** | $23.20 \pm 0.94$ | $20.87 \pm 2.10$ | $21.83 \pm 0.91$ |
| **β-sheet (%)** | $28.50 \pm 0.87$ | $31.07 \pm 2.40$ | $29.1 \pm 2.78$ |
| **β-turns (%)** | $11.97 \pm 0.08$ | $12.43 \pm 0.84$ | $12.03 \pm 0.40$ |
| **Unordered (%)** | $36.34 \pm 0.59$ | $35.63 \pm 1.02$ | $37.03 \pm 2.61$ |

<sup>a,b,c,d</sup> indicates statistical difference ( $p < 0.05$ ) using one-way ANOVA between cLF, pfLF\_highFe and pfLF\_lowFe obtained by CD and FTIR measurements.

For all preparations, unordered regions were observed to be the predominant secondary structure element, followed by β-sheets, α-helices and lastly β-turns, again consistent with existing literature (Fang et al., 2014; Mizutani, Toyoda, & Mikami, 2012). These findings are also consistent with prior research showing only small changes in secondary structure as a function of iron saturation (Sreeedhara et al., 2010; Wang et al., 2013; Xia et al., 2023). Despite the similarities in secondary structure, there does appear to be a general observable trend depending on the protein’s iron saturation. Specifically, we observe a slight increase in α-helix and a slight decrease in β-sheet percentages as the amount of bound iron decreases. As previously mentioned, it is well understood that decreasing iron saturation results in a more open, flexible conformation of the lactoferrin molecule (Baker & Baker, 2004), which may favour the formation of α-helix within the protein. Statistical analysis revealed a significant difference between α-helix content of pfLF_highFe when compared to pfLF_lowFe and cLF, but of no other secondary structure component, suggesting that a very good structural conservation can be achieved *via* precision fermentation.

CD was also employed to further validate secondary structure estimations (**Figure 5b**). The CD spectra of all preparations display a similar trajectory, beginning with a maximum positive mean residue ellipticity (∼4 degrees cm² dmolc¹) at 190 nm, attributed to α-helical structures. The spectra sharply decrease, crossing zero at ∼203 nm, and descend to a significant negative trough near 208 nm, indicative of β-sheets. Beyond 208 nm, ellipticity decreases less steeply, reaching a second, less pronounced minimum at 222 nm, before leveling off towards zero as the wavelength approaches 260 nm.

Secondary structure contributions were quantified using the CONTINN algorithm *via* Dichroweb. Results, presented alongside FTIR data in **Table 2**, confirm dominance of unordered regions, followed by β-sheets, α-helices and β-turns. The observed trends and relative structural contributions are consistent with the findings of FTIR analysis. Statistical analysis revealed no significant differences across groups. Overall, it appears that iron saturation and the different drying techniques applied to PF and commercial bLF have minimal impact on lactoferrin’s overall secondary structure, with a strong structural conservation again being observed through precision fermentation.

### 3.7 Thermal Stability

The thermal denaturation profiles of lactoferrin preparations were assessed using DSC at a scan rate of 3°C/min, with thermograms shown in **Figure 6**. It appears the thermal denaturation properties of each variant are influenced by iron saturation. pfLF_highFe, with 96.26% iron saturation, exhibits a single denaturation peak, whereas preparations with more native iron levels, pfLF_lowFe and cLF, display two distinct peaks (**Table 3**).

**Fig. 6.**
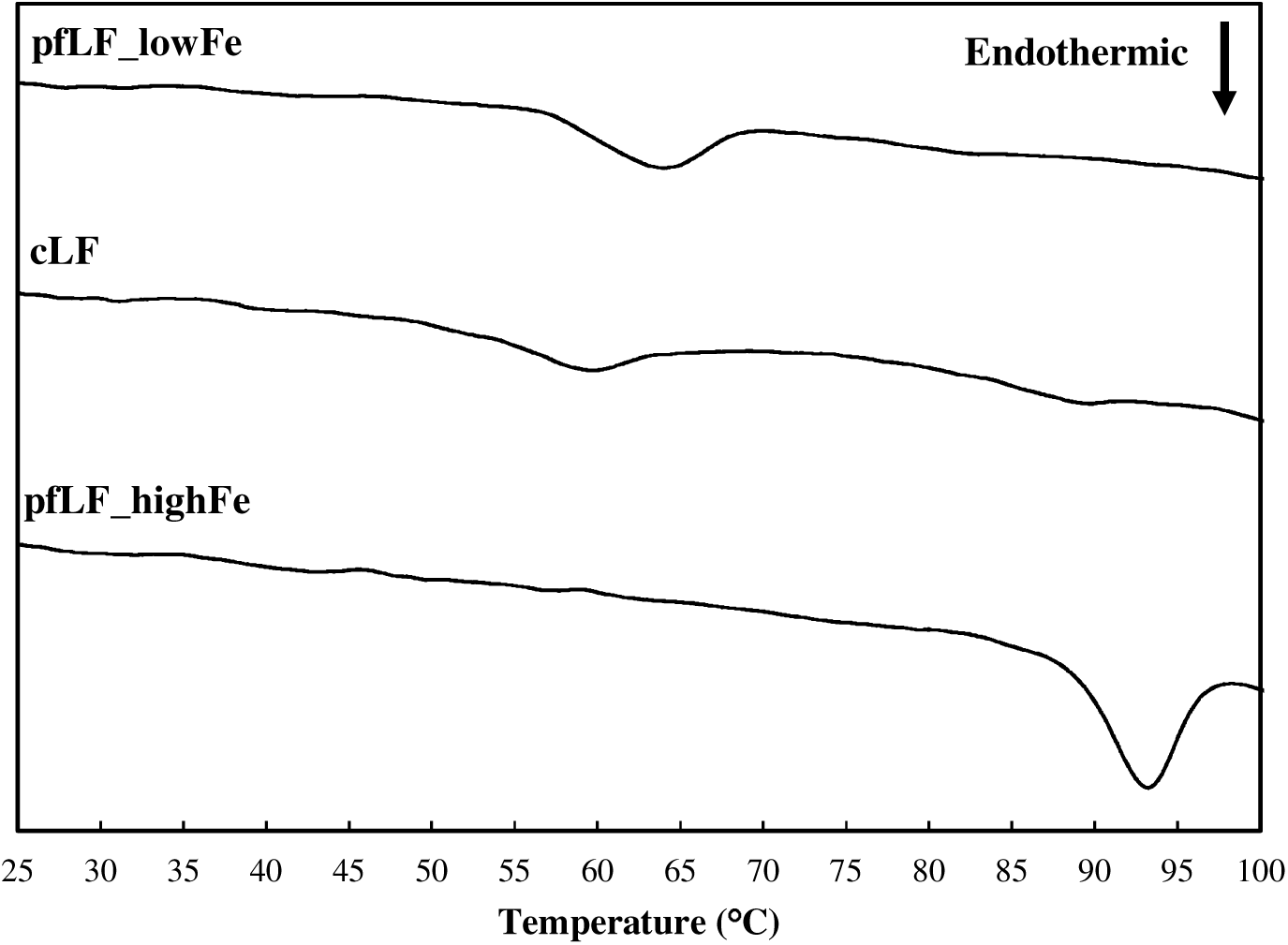
DSC thermograms of 10 mg/mL cLF, pfLF_highFe and pfLF_lowFe at a scan rate of 3°C/min.

**Table 3.** Thermal stability of 10% (w/w) cLF, pfLF_highFe and pfLF_lowFe at a scan rate of 3°C/min.

| Sample | T<sub>s</sub>1 (°C) | T<sub>m</sub>1 (°C) | T<sub>s</sub>2 (°C) | T<sub>m</sub>2 (°C) | ΔH (J/g) |
| --- | --- | --- | --- | --- | --- |
| cLF | 51.97 ± 0.74 | 59.19 ± 0.06 | 83.82 ± 0.03 | 88.40 ± 0.37 | 1.11 ± 0.04 |
| pfLF_lowFe | 57.19 ± 0.08 | 63.50 ± 0.34 | 76.11 ± 0.83 | 82.88 ± 0.23 | 1.28 ± 0.04 |
| pfLF_highFe | 88.52 ± 0.04 | 93.15 ± 0.05 | N/A | N/A | 3.62 ± 0.44 |

This behaviour may reflect lobe-specific thermal stability, with Anderson et al. (1990) demonstrating *via* X-ray crystallography that iron binding causes the C-lobe to become more compact than the N-lobe, possibly leading to distinct denaturation events. Alternatively, the presence of monoferric lactoferrin molecules may explain the observed thermograms. This ‘monoferric’ hypothesis postulates that the major peak in native-LF preparations is the result of monoferric lactoferrin, whereas the minor peak may be the result of relatively low proportions of diferric lactoferrin molecules (Rüegg, Moor, & Blanc, 1977).

An apparent increase in T_m1_ (59.19 ± 0.06°C → 63.50 ± 0.34°C) and decrease in T_m2_ (88.40°C ± 0.37 → 82.88 ± 0.23°C) for pfLF_lowFe when compared to the cLF sample, which has a slightly higher iron saturation is observed. This is likely due to an increased proportion of apo-LF in the lower iron sample, shifting T_m_ values towards the expected value of 70°C for apo-LF (Bokkhim et al., 2013; Paulsson et al., 1993). In contrast, holo-LF (pfLF_highFe), likely exists in an almost entirely diferric state, hence exhibiting only a single T_m_ peak at 93.15 ± 0.05°C, with negligible monoferric populations available to produce earlier peaks.

Irrespective of the underlying physical process, the observed thermal behaviours align with previously established findings on LF thermal stability as a function of bound iron (Bengoechea et al., 2011; Bokkhim et al., 2013; Paulsson et al., 1993), and again highlight the efficacy of structural conservation through precision fermentation.

### 3.8 Surface Hydrophobicity

Surface hydrophobicity measurements are important to understand the functionality and application of proteins in formulations as findings may correlate with a protein’s emulsification capability (Schofield, 1997). Globular proteins minimise the energetic cost of exposing hydrophobic residues to water by burying them in their core, resulting in a stable, energetically favourable conformation (Sun, 2022). As such, measurements of surface hydrophobicity may also allow differences in protein structural assembly to be probed.

**Figure 7** details the Surface Hydrophobicity Index (S_0_) of the three lactoferrin preparations, with findings demonstrating that cLF recorded a notably high S_0_ value of approximately 390, when compared to the much lower values produced by pfLF_highFe and pfLF_lowFe (21.20 and 81.75, respectively). Results for the PF samples again align with previous reports of lactoferrin surface hydrophobicity in which an increase in S_0_ is observed as iron saturation decreases, or the structure becomes more open (Goulding et al., 2021). The significantly higher S_0_ of cLF compared to pfLF_lowFe, despite similar iron saturation, is likely due to spray drying-induced partial denaturation, which exposes hydrophobic residues that interact more readily with the ANS probe. This is consistent with findings in existing literature, with Liu et al. (2016) observing that S_0_ values for a native bLF preparation increased by an order of magnitude following a heat treatment (90°C for 10 minutes). Interestingly, the observed denatured state of the commercial sample did not directly translate to any dramatic alterations in secondary structure (as confirmed by FTIR and CD analysis). This finding, however, is also consistent with literature, which reports that thermal treatment of lactoferrin can preserve secondary structure composition, while still forming partially folding intermediates with a more disordered tertiary structure reflecting a molten globule state (Schwarcz et al., 2008). Whether these structural changes manifest as functional differences is explored in the subsequent section.

**Fig. 7.**
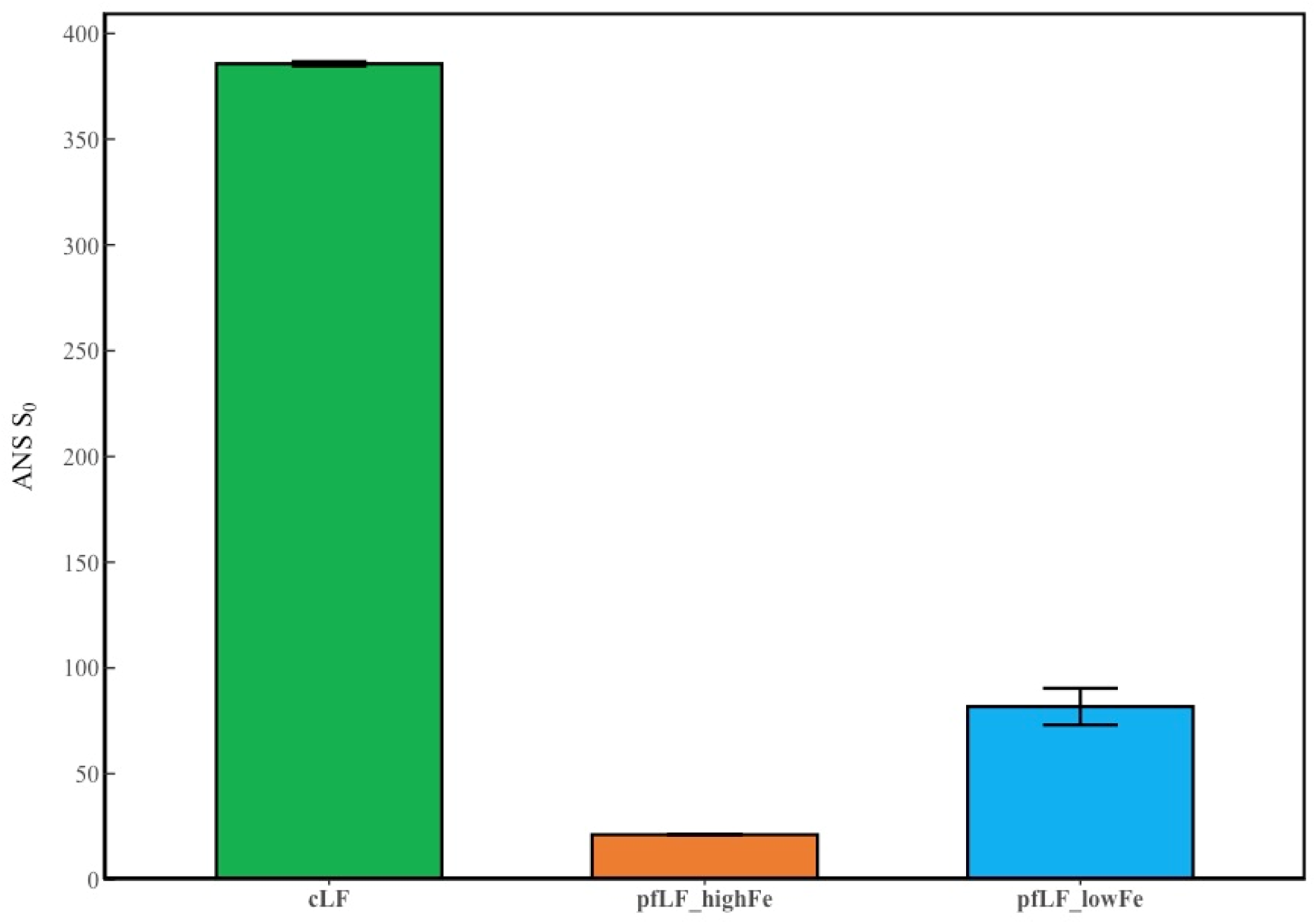
Surface hydrophobicity index (S_0_) of cLF, pfLF_highFe and pfLF_lowFe using ANS fluorescent probe binding assay.

### 3.9 Antimicrobial and Anti-inflammatory properties

The inherent ability of iron to donate or accept electrons in cellular redox processes highlights the crucial role it plays in supporting both the survival and functional integrity of many organisms (Mazurier & Spik, 1980; Parrow, Fleming, & Minnick, 2013). This role emphasises the importance of iron-chelating proteins such as lactoferrin which can sequester free iron within mammalian hosts and restrict bacterial growth. An adapted antimicrobial assay described by Sekse et al. (2012) was employed to probe the inherent antimicrobial properties of each lactoferrin preparation (**Figure 8a**). pfLF_lowFe and cLF exhibited very similar (0.187 mg/mL and 0.170 mg/mL) minimum inhibitory concentrations (MIC) against *E. coli* W3310 (K12) compared to pfLF_highFe, which showed no growth inhibition. This result is therefore attributed directly to the iron saturation and hence iron binding capacity of the lactoferrin preparations given the two pfLF proteins are identical apart from the different iron saturation status.

**Fig. 8.**
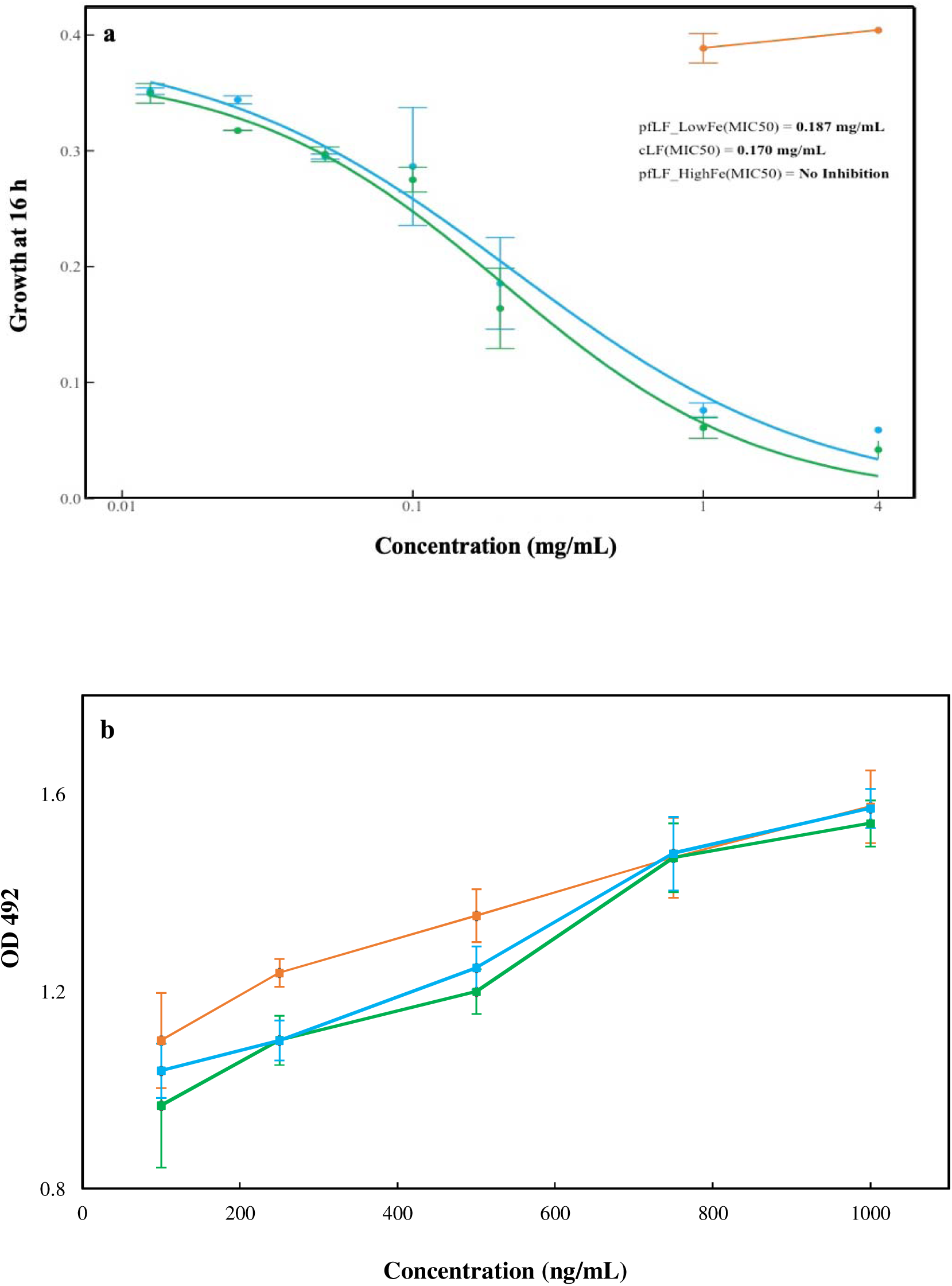
(a) Growth of *E. coli* strain W3110 (K12) at 16 hours as a function of lactoferrin concentration (mg/mL) for cLF (green), pfLF_highFe (orange) and pfLF_lowFe (blue). Growth inhibition is quantified as the concentration required to achieve a 50% reduction in growth (MIC50). (b) Binding of cLF (green), pfLF_highFe (orange) and pfLF_highFe (blue) to Lipid A*. E.coli,* natural biphosphoryl as measured by absorbance at OD 492nm.

While iron chelation is an important functional property of lactoferrin, it is now understood that direct binding of the protein and its peptides to microbial surfaces further enhances its broad-spectrum bioactivity. Within the N1 subdomain of bLF, an 18-loop region with high positive charge density and hydrophobicity exists that allows the protein to interact strongly with structural components of bacteria which are anionic in nature. Specifically, it appears that a 6-residue region, ^20^RRWQWR^25^, is critical in this interaction due to the positive charge carried by arginine (R) and the hydrophobicity of tryptophan (W), which can aid destabilisation and increased permeability of bacterial membranes through hydrophobic interactions (Drage-Serrano et al., 2012). This interaction also appears to play an anti-inflammatory role by limiting the presentation of bacterial lipopolysaccharide (LPS) to Toll-like receptor 4 (TLR4), reducing the production of inflammatory cytokines and improving protection against bacterial infections (Ling & Schryvers, 2006). Analysis detailed in **Section 3.1** showed that the amino acid sequences remained consistent across the three preparations and they each contain the critical 18-loop region that should facilitate bLf-LPS interaction. It was therefore expected that each preparation would show an interaction with Lipid A diphosphoryl from *Escherichia coli* F583 (Rd mutant) and results for this analysis are shown in **Figure 8b**. Each sample showed increased binding with LPS as a function of increasing concentration, reflected by increasing OD at 492 nm. It may be inferred from these interactions that cLF, pfLF_highFe and pfLF_lowFe all share the antibacterial and anti-inflammatory therapeutic potential that has been previously described (Ellison, Giehl, & LaForce, 1988; Yamamuchi et al., 1993; Döhler & Neberman, 2002).

The control containing only lactoferrin (data not shown) showed no increase in absorbance at 492 nm as the concentration increased. This indicates that the observed results arise from interactions with LPS rather than artifacts from direct lactoferrin binding to the ELISA plate. Despite the differences in iron saturation experienced by each preparation, no clear link between iron saturation % and LPS binding capacity seems present. These findings further emphasise that iron chelation and sequestration are not the sole driving force behind lactoferrin’s functional capabilities. Importantly, findings also highlight that not only can precision fermentation produce proteins with exact primary structure fidelity, but that this structural conservation is mirrored in the preservation of bio-functional properties.

## Conclusions

Precision fermentation is demonstrated to produce bLF that competes with current commercial offerings. PF samples are shown to have a comparable primary sequence coverage to their commercial counterpart and statistically identical secondary structural components (at similar iron levels). Similarly, molar absorption and pI of the samples were within the same range and differences in tertiary structure, thermal denaturation profile and iron chelation ability were attributed to varying iron saturation levels. Crucially, the bio-functional properties of the PF samples closely mimic those of the commercial sample at equivalent iron levels. Notably, the PF method can produce the protein in its native conformation, to the desired level of iron saturation with minimal thermal intervention, preserving its native surface hydrophobicity. The successful production of bLF *via* PF has broad implications for the development of non-animal sourced functional foods and therapeutics, and future studies should aim to characterise the techno-functional application of precision-fermented bLF with comparison to its commercial counterpart.

